# Baffling: A cheater strategy using self-made tools in tree crickets

**DOI:** 10.1101/2020.05.06.080143

**Authors:** Rittik Deb, Sambita Modak, Rohini Balakrishnan

**Affiliations:** Centre for Ecological Sciences, Indian Institute of Science, Bangalore, India

## Abstract

Intense sexual selection in the form of mate choice can facilitate the evolution of different alternative reproductive strategies. These strategies can be condition-dependent, wherein genetically similar individuals express the strategy based on their condition. Our study shows that baffling, a mate attraction strategy using self-made acoustic amplifiers, employed by male tree crickets, is a classic example of a condition-dependent alternative strategy. We show that less preferred males, who are smaller or produce less loud calls, predominantly use this alternative strategy. Baffling allows these males to increase their call loudness and advertisement range, and attract a higher number of mates. Baffling also allows these males to deceive females into mating for longer durations with them. Our results suggest that the advantage of baffling in terms of sperm transfer is primarily limited to less preferred males, thus maintaining the polymorphism of calling strategies in the population.

**Impact statement:** This study shows that less preferred tree cricket males use an alternative signaling strategy to call louder, thus attracting and mating with otherwise choosy females using deception.

## Background of the work

Sexual selection is a dominant force that generates significant variation in biological systems. This variation is not, however, limited to diversity between the sexes. Sexual selection operating in tandem with natural selection can generate significant diversity within each sex. One of the primary examples of within-sex diversity is alternative reproductive strategies and tactics. Taborsky et al. (2008) summarised these as the presence of discontinuous phenotypic traits (physiology, morphology, behaviour) that maximize individual fitness without using the dominant strategy. Generally, such strategies become frequent when intense sexual selection creates biased mating and skewed reproductive success in a population. In nature, the strength of operational sexual selection being stronger on males has led to the prevalence of such tactics predominantly in this sex (Shuster, 2010; Taborsky et al., 2008).

In the tree cricket genus *Oecanthus*, males attract females using acoustic signals: they primarily call from leaf edges, and females respond by localizing the males (Walker, 1957). Interestingly, there exists an alternate mode of signaling, where the males make a hole at the center of a leaf and use it as an acoustic baffle (a sound amplifier) (Prozesky-Schulze et al., 1975). The use of this strategy increases the loudness of the call by reducing acoustic short-circuiting (Forrest, 1982; Prozesky-Schulze et al., 1975). Our previous work (Mhatre et al., 2017) showed that male tree crickets not only manufacture these tools (Mhatre, 2018) but do so optimally: they choose larger leaves and make holes in optimal leaf positions, maximising sound amplification. However, to date, it remains unclear what drives the evolution and maintenance of this alternative behavioural strategy in the population.

In this study, using extensive field sampling and laboratory experiments, we show that the baffling propensity is higher in smaller and softer males (the term ‘softer’ refers to lower call intensity). Interestingly male body size is known to be under strong selection pressure from female mate choice: females mate longer with larger males (Deb et al., 2012), leading to transfer of higher number of their sperms compared to smaller males. Using behavioural experiments and simulations, we show that baffling is a successful cheater strategy that is used by less preferred males to gain mating and reproductive benefits. Our results suggest that the advantage of this strategy is mostly limited to less preferred males, making it a classic example of a condition-dependent alternative strategy.

## Results

### Who are the bafflers?

Our field sampling (Fig S1) revealed that baffling is rare (25 bafflers out of 463 calling males sampled (5%)) in the natural condition. Our earlier study (Mhatre et al., 2017) had shown that not all males prefer to call from baffles even when given ideal leaf sizes. Our field observations revealed that males calling from baffles (henceforth termed as bafflers) were smaller in body size than non-baffling males (henceforth termed as non-bafflers) (Welch t = 4.09, df = 42.65, *P* = 0.0002) (Fig. 1A). We also found that bafflers when calling without a baffle, had lower sound pressure levels (SPL: a measure of call loudness) than non-bafflers (Welch t =2.93, df = 24.98, *P*= 0.007) (Fig. 1B). Our controlled laboratory experiments showed that propensity to baffle was higher on larger leaves (Analysis of deviance: χ^2^_3_=8.63, *P* = 0.03, Table S1B) (concordant with our previous study Mhatre et al. (2017)), and for smaller (Analysis of deviance: χ^2^_2_=8.08, *P* = 0.02, Table S1B) and softer males (i.e. males with lower SPL) (Analysis of deviance: χ^2^_1_ = 10.42, *P* =0.001, Table S1B) (Fig 1C and D) (see methods section for experimental details). Smaller males baffled with a higher propensity, which increased with increase in leaf size (extra-large leaf: medium & small males vs large males: Proportion test χ^2^ = 5.95, df = 1, *P* = 0.02, large leaf: small vs medium males: Proportion test χ^2^ = 5.95, df = 1, *P* = 0.02, large vs medium males *P* = 0.46) (Fig.1C). For smaller leaves, the propensity to baffle was not influenced by male size (medium leaf: *P* = 0.91 and small leaf: *P* = 0.94) (Fig.1C) owing to inherent lack of baffling using these leaf sizes (as also shown in Mhatre et al., 2017). Bafflers when calling without baffles had lower SPL calls (softer calls) than non-bafflers, independent of leaf size (small leaf: sample size too low for statistical testing, Medium leaf: t = -8.63, df = 11.67, *P* =2.09×10^−6^, Large leaf: t = -10.46, df = 46.1, *P* =9.29×10^−14^, Extra-large leaf: t = -6.19,df = 28.2, *P* =1.07×10^−6^) (Fig. 1D).

**Fig 1:**
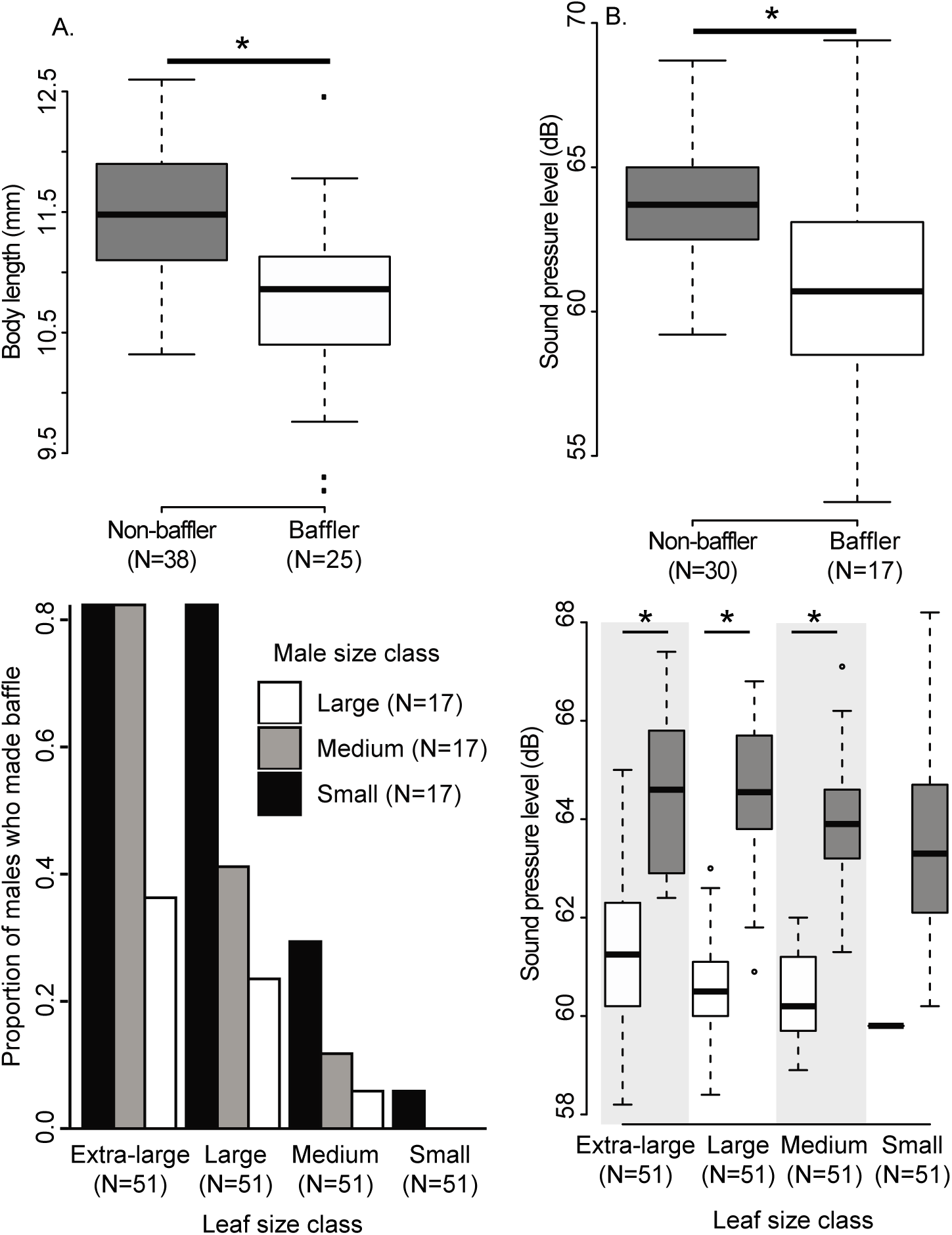
Relation between baffling propensity, male size, and call SPL. A) Box plots showing the body size distribution of bafflers and non-bafflers observed in the field. The ‘*’ indicates a significant difference. B) Box plots showing the without baffle call SPL distribution of bafflers (when forced to call from leaf edge), and non-bafflers in the field. The ‘*’ indicates a significant difference. C) Bar diagram showing baffling propensity as a function of leaf size and male body size. Predominantly small males were the bafflers across leaf size classes. Leaf size classes: small (length: 35 ± 2 mm), medium (length: 50 ± 2 mm), large (length: 65 ± 2 mm), and extra-large (length: 115 ± 4 mm). Male size classes: small (length: <10.9 mm), medium (length: 10.9-12.08 mm), large (length: >12.08 mm). This was a repeated measures design where the same 17 males of each size class; i.e., 51 males were tested on each leaf size. D) Box plots showing the distributions of call SPLs of non-bafflers (grey boxes) and bafflers when calling without baffles (white boxes) across three different leaf size classes. The ‘*’ indicates a significant difference. Please note missing ‘*’ symbol for small leaf, as the number of baffler in small leaf was only 1; hence no test was performed. However, the pattern was comparable to other leaves.

### What are the advantages of baffling?

We know that baffling increases call SPL (call loudness) (Forrest, 1982; Mhatre et al., 2017; Prozesky-Schulze et al., 1975); however, its consequences for individual fitness are unknown.

### Increase in active acoustic volume

We found that by baffling, males gained an increase in SPL (loudness) between 8-12 dB (mean ± SD: 10.2 ± 1.2 dB, Welch t-test=35.4, df =16, *P* < 2.2×10^−16^) in the wild (Fig. S2A). Our previous work (Mhatre et al., 2017) indicated that sound radiation efficiency is maximized in large leaves. We found comparable results in empirical measurements of gain in SPL where larger leaves resulted in higher SPL gain (loudness gain) (Fig. S2B, Table S2). We also found that gain in SPL (loudness) was higher for smaller males, especially when calling from small leaves (Fig S2B, Table S2). We calculated the shape and volume of 10 free moving males’ active acoustic volumes in the laboratory using a customized set-up (Fig. S3A) while calling from both leaf edge and a baffle. The active acoustic volume of a calling male is defined as the volume within which the calling SPL is greater or equal to the behaviorual hearing threshold of the female. The acoustic volume resembled a 3-dimensional figure of ‘8’ (Fig S3B, C, and D) with the animal at the center. This shape was well approximated (R^2^ = 0.99, *P* <2.2×10^−16^) (Fig. S4B) by two identical ellipsoids touching each other along the longest radius (b) representing the body axis of the animal (Fig S4A). We calculated active acoustic volumes of males calling at different SPLs (loudness) (ranging from 58-68 dB) with varying baffling gain (ranging from 8-12 dB). We found that baffling caused substantial increment (2.5-11 times) of the overall active acoustic volume (Fig 2A).

**Fig 2.**
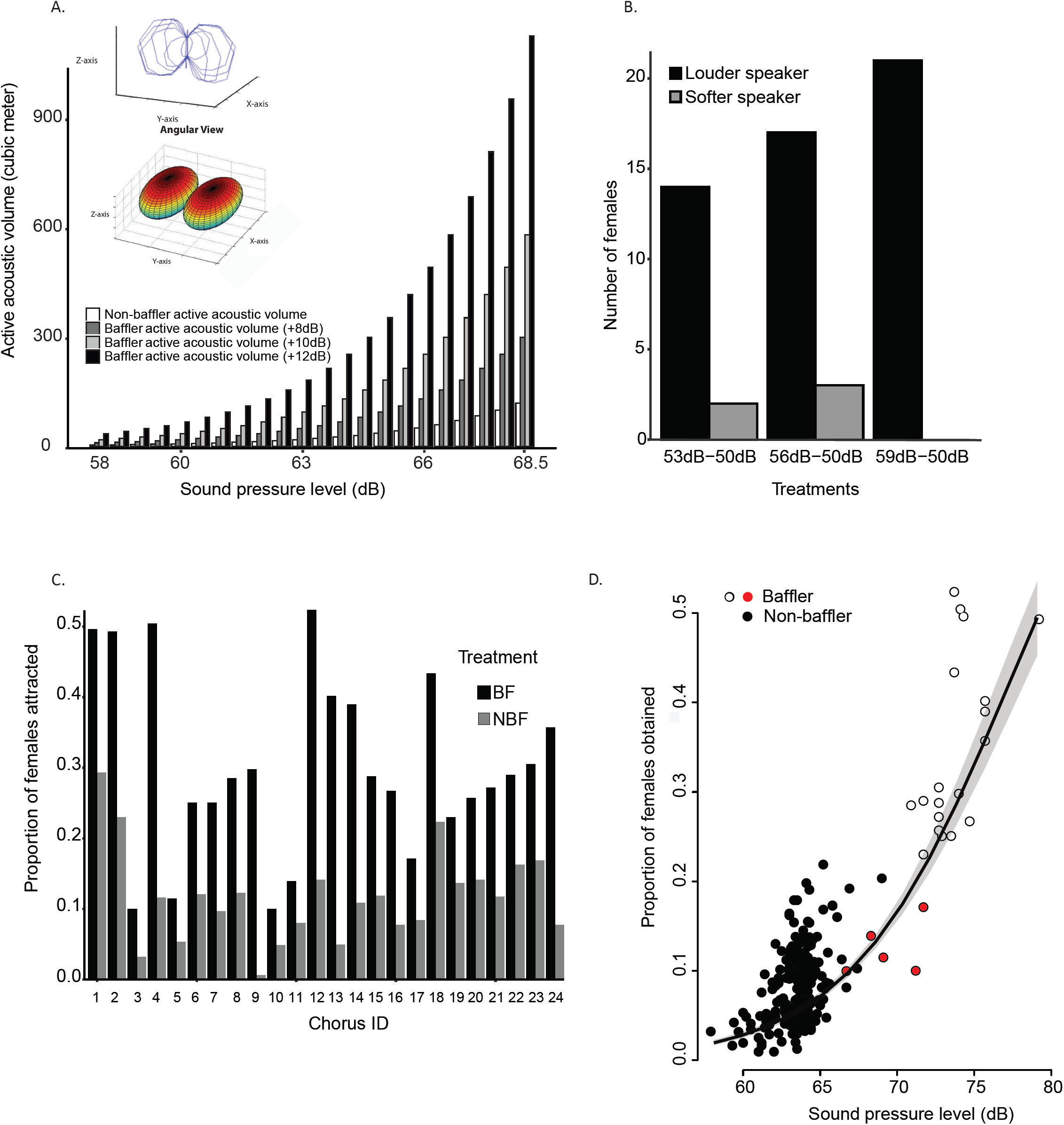
Call SPL, female preference, and baffling. A) A bar plot showing the comparison between the active acoustic volume of an individual when calling without a baffle (Non-baffler volume) and when it is baffling with either 8, 10 or 12 dB increment in call SPL. The plot shows this across a range of non-baffling call SPLs from 58 to 68.5 dB (x-axis). This plot clearly shows that baffling caused a significant increase in active space volumes. Inset showing wire-diagram and approximate shape of the active acoustic volume. B) Bar plot showing female response in the two speaker choice experiment. The females showed a preference for the louder call (black bars) in all the three treatments. The sample size for 50-53 dB, 50-56 dB and 50-59 dB treatments were 16, 20 and 21 females respectively. C) A bar plot showing the difference in the proportion of females attracted by a single male (same individual) when it was baffling (black bar) vs. when it was not baffling (grey bar) across 24 simulated natural choruses. D) A scatter plot showing the relationship of calling SPL and the proportion of females obtained by the caller (pooled across all the animals in 24 choruses for 100 iterations in each chorus) when one individual in each chorus was transformed into a baffler (Scenario 2). White or red circles represent the bafflers. The red circles denote the bafflers who did not obtain the highest proportion of females in their respective chorus despite baffling. The line shows the fit of the GLM model along with 95% CI (the grey area around the trend line).

### Female phonotactic preference for louder calls

Using the natural male chorus structures (Deb and Balakrishnan, 2014), we calculated the SPL (loudness) differences that females predominantly encounter in the field (50 dB median, with difference of 9dB) (see methods, Fig S5A). Our two choice phonotactic assay (Fig S5B) revealed that females preferred to approach the louder calls, even when the difference in SPL was just 3 dB (50:53 dB: χ2 = 9, df = 1, *adj.P* = 0.003, 50-56 dB: χ2 = 9.8, df = 1, *adj.P* = 0.002, 50-59 dB: χ2 = 21, df = 1, *adj.P*= 4.59×10^−6^, Fig 2B).

### Increase in the proportion of females attracted

We combined the results of active space volume overlap in the field and female preference for louder calls to simulate 24 natural choruses (Deb and Balakrishnan, 2014) to examine whether bafflers attracted greater proportion of females. We calculated this by estimating the number of females obtained by each male in a chorus after calling for 100 nights, and dividing it by the total number of females in a chorus across 100 nights (considering 100 nights as life expectancy and sex ratio 1:1, see methods). In the choruses where none of the males were baffling (scenario 1, see methods), louder the male, higher was the proportion of females obtained (Fig S6A, calling SPL: 0.18, std.error = 0.02, z = 8.35, *P* = 6.54 X 10^−15^). Next, we converted a randomly chosen male in each chorus into a baffler by increasing its call SPL (loudness) and changing the active acoustic volume (scenario 2). We found that the proportion of females attracted increased significantly during baffling (Fig. 2C) in each chorus (Pairwise-Permutation test: BF - NBF = 4.643, *adj.P* = 3.43 X10^−06^). This was interesting, as our simulation allowed a non-baffling male to rotate and advertise around 360° axes within a night (a natural phenomenon), whereas bafflers’ advertisement directions were restricted (defined by the angle of the chosen leaf). Despite the lack of omnidirectional signaling, the bafflers gained in the proportion of females attracted. We found that the proportion of females attracted by the bafflers increased with increasing SPL (Fig 2D, calling SPL: 0.16, std.error= 0.007, z = 24.03, *P* = 2×10^− 16^, see Fig. S6B for Z-score plot). Baffling allowed males to obtain the highest proportion of females in 19 out of 24 choruses (Fig 2D, white circles).

### Do females get deceived by bafflers?

Our earlier work showed that tree cricket females prefer larger males during mating, even in the absence of acoustic cues (Deb et al., 2012). This preference was manifested by retaining the spermatophores of the preferred males significantly longer (Deb et al., 2012), thus allowing longer duration for sperm transfer (Brown, 1997). This posed the interesting question, whether females could differentiate between bafflers (smaller/softer call males) and genuine louder/ bigger callers, during mating? This experiment was crucial to understand if baffling could effectively work as a cheater strategy by deceiving females. A playback and mating experiment was designed to replicate a natural phonotaxis followed by mating scenario (see methods). Each night, either a soft (< 61 dB SPL) or loud (> 66 dB SPL) caller male was caught from the field (as SPL has high variability across nights, Deb et al. (2012)). For the playback experiment, we chose a total of 20 soft callers and increased their call SPL by 8-12 dB via playback. This effectively transformed them into bafflers from a female perspective (SL: soft transformed to loud) We kept the call SPLs of another 20 soft males unaltered during playback (SS: soft remained soft). Similarly, we designed playback treatments for naturally loud callers (10 males, LL: loud remained loud, and 10 males, LEL: loud transformed to extra loud). If females could identify the genuine loud callers, we expected that they would discriminate between LL and SL males and would not discriminate between SL and SS males. Whereas, if baffling influenced female choice during mating, we expected females to mate longer with the SL males as compared with SS males, and to not discriminate between SL and LL males. We had two treatment effects: a) body size of male (small, medium or large) and b) playback treatment (SS, SL, LL, and LEL) resulting in 12 treatment combinations (Fig 3).

**Figure. 3.**
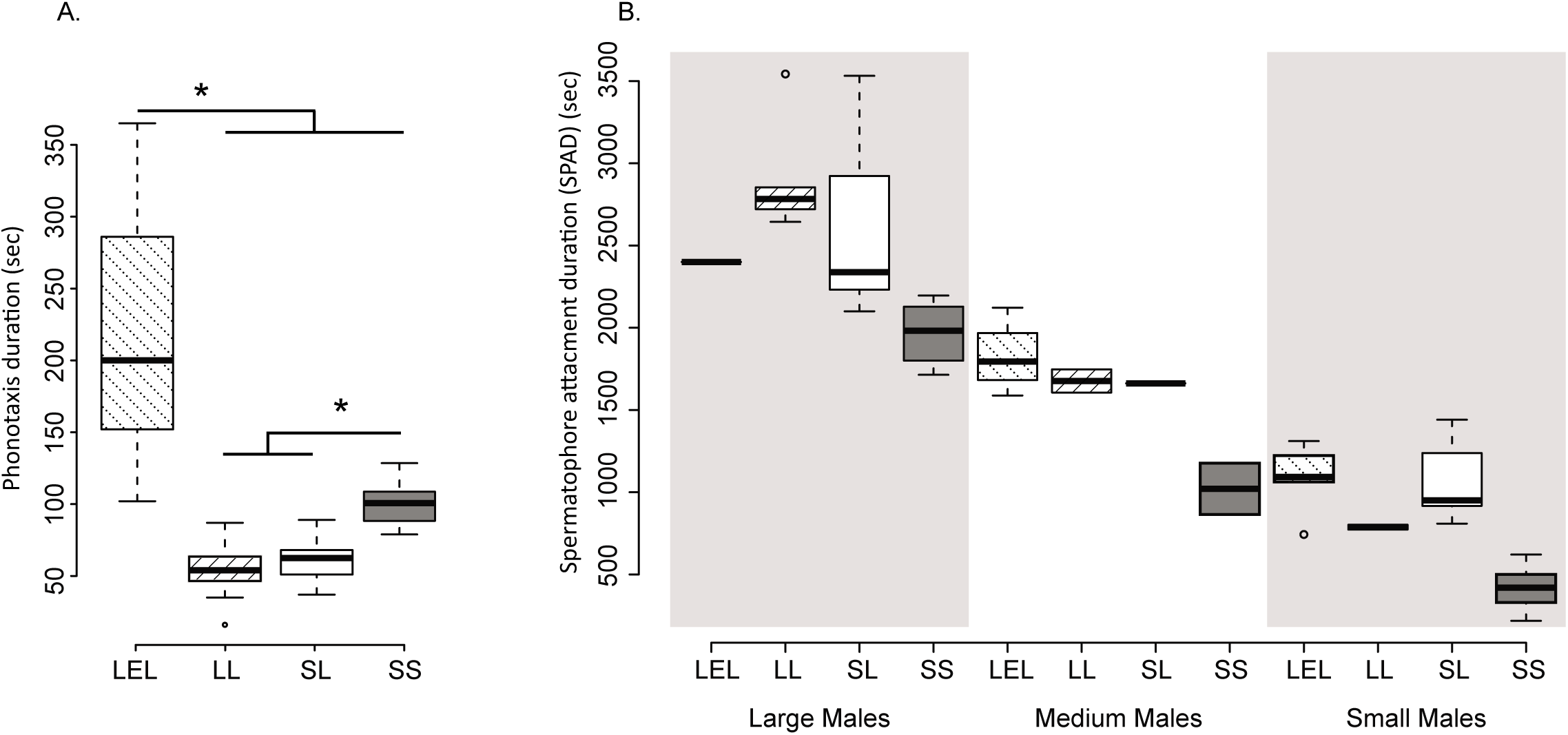
The benefits of baffling for small and large males. A) Box plots showing phonotaxis duration across 4 playback treatments. Abbreviations: SS (grey box): soft caller whose call was played back as a soft call (n=20), SL (white box): soft caller whose call was played back as a loud call (n=10), simulating baffling, LL (right-angled striped box): loud caller whose call was played back as a loud call (n=20), LEL (left-angled striped box): loud caller whose call was played back as an extra loud call, simulating baffling (n=10). The ‘*’ indicates a significant difference. B) Boxplots showing the distributions of spermatophore attachment duration (SPAD) across different treatments. SPAD is a good indicator of sperm transfer time, which is often related to male reproductive success. These treatments were carried out across different body-sized males; Male size classes: small (length: <10.9 mm) (26 males), medium (length: 10.9-12.08 mm) (9 males), large (length: >12.08 mm) (25 males).

We found that the female phonotactic response varied depending on the playback treatment (F_3,56_= 47.5, *P* =2.06×10^−15^). The females did not differentiate between a true loud caller (LL) and a baffler (SL) in the time taken to approach the call during phonotaxis (SL vs. LL, estimate: 7.85, t = 0.56, *P* = 0.9) (Fig 3A). However, they showed a faster response towards louder calls (SS vs SL, estimate: 38.5, t=2.74, P=0.04, SS vs LL, estimate: 46.35, t=4.04, *P* < 0.001) (Fig 3A). Interestingly, we found that the females corrected their path multiple times and took significantly longer to reach the LEL calls(LL vs. LEL: estimate - 160.35, t = -11.4, *P* < 0.01, SL vs. LEL: estimate -152.5, t =-9.4, *P* < 0.01, SS vs. LEL: estimate -114, t =- 8.1, *P* < 0.01). We speculate that this response was due to its loudness (75-80 dB) being close to auditory saturation for the female, thus creating localization errors.

We found that spermatophore attachment duration (SPAD) varied depending on both playback treatment and male body size (playback treatment: F_3,54_ = 42.02, *P* =3.79×10^−14^, Male size class: F_2,54_=181.88, *P*=2×10^−16^) (Fig 3B). We found that the SPAD was shorter for softer (SS) males in all size classes (LEL vs. SS: -695.8, t =-5.95, *P* = 2.03×10^−7^) (Fig.3B) indicating female discrimination against soft callers during mating. The SPAD was comparable between SL, LL and LEL males (LEL vs. SL: -58.5, t = -0.49, *P* = 0.62, LEL vs. LL: 47.9, t = 0.35, *P* = 0.72) (Fig 3B) indicating the efficacy of baffling in deceiving females into mating for longer durations with the less preferred softer males than they otherwise would have. Concordant with our previous study (Deb et al., 2012), SPAD increased with male body size (large vs small: -1589.8, t = -19.07, *P* = 2×10^−16^, large vs medium: -937.5, t = -7.79, *P* = 2.12×10^−10^, Fig 3B). However, it was evident that the differences in SPAD between larger and smaller males declined when the smaller males baffled (Fig 3B). These results mean that a baffler can deceive a female into approaching and mating with it for longer durations than if it did not baffle. Baffling thus provides a two-fold advantage: increasing the likelihood of female approach and increasing SPAD during mating, which in the genus *Oecanthus* is positively correlated with the number of sperms transferred to the female (Brown, 1997).

### Final puzzle: why don’t larger and louder males baffle as much?

We hypothesized that the reason why larger and louder males did not baffle as frequently, was because they gained no/little/variable benefits from baffling. We expect that baffling gain would be limited by a) the maximum number of matings possible per night (activity duration divided by mating duration), and b) the number of sperms transferred. As louder and larger males attracted a significant number of mates/night (mean ± sd: 2.31 ± 0.7) and had very high SPAD (∼45 minutes/female) (Deb et al., 2012) even without baffling, we expected their gain by baffling to be minimal.

Using the data from our previous experiment (Fig 3B), we compared the gain in SPAD by baffling for soft males (i.e., SL vs. SS) across two body size classes (large and small). Primarily we compared if the increase in SPAD for a small and soft baffler (calculated as the difference between S_SL and S_SS) was higher than the increase in SPAD for a large and soft baffler (calculated as the difference between L_SL and L_SS) (Fig.3B) (see methods). We found that small males (median gain: 624 sec) gained significantly longer SPAD than large males (median gain: 483.5 sec) by baffling (W = 4252.5, *P* = 0.04).

We finally used simulations to compare the advantages of baffling (both numbers of females attracted and sperm transferred) between different classes of males (loud vs. soft, and small vs. large) across 23 (22 for SPAD) natural choruses. Our simulation incorporated animal activity duration (120-140 min/night), and mating duration (large males: ∼42 mins, small males: ∼15 mins (see Fig 3B)) obtained from our earlier studies on this species (Deb and Balakrishnan, 2014; Deb et al., 2012). We used the rate of sperm transfer (sperm transfer function) of *O. nigricornis* (a congeneric species to *O. henryi*) to calculate the total number of sperms present in the spermatophore of a male and used it for the simulation (Brown, 1997) (see methods). Our simulation used a nested design. In 22 natural choruses, we randomly chose (using a random number generator in MATLAB) a loud male and examined the proportion of females it attracted when it was baffling and when not baffling. For each of these loud baffling males, we considered two scenarios: that it was a a) large, and b) small male. We calculated the number of mates attracted and estimated the number of sperm transferred during mating (see methods) in both scenarios. Our simulation allowed re-mating within a night, which was limited by the amount of time spent on individual mating and the total activity period observed in this species. We kept the activity period comparable between a large and a small male, which allowed maximum 2 matings for a large male (a total of 84 minutes) and up to 5 matings for a small male (a total of 75 minutes). We pooled the number of sperms transferred by each male within a night. We simulated each chorus 100 times (corresponding to 100 nights) to calculate a male’s lifetime sperm transfer (LST) success. We calculated the difference of LST for a loud, large male when it was baffling and when it was not baffling, and repeated the same analysis for loud, small males. We repeated this entire simulation for soft_large and soft_small males and finally compared the distributions to understand the gain differences.

We found that soft non-baffler males, soft_baffler males, loud non-baffler males, and loud_baffler males obtained a median of 0.35, 2.5, 2.6, and 7.7 females/night respectively (Fig 4A). However, as a male (soft or loud) could mate with only a limited number of females/night (calculated as activity period divided by mating duration) its mating opportunity was limited (Fig 4A). Our analysis clearly showed that it was only advantageous for a soft male to baffle, as the louder males were attracting close to the maximum number of females they could mate with within a night (around 3) even without baffling (Fig 4A).

**Fig. 4.**
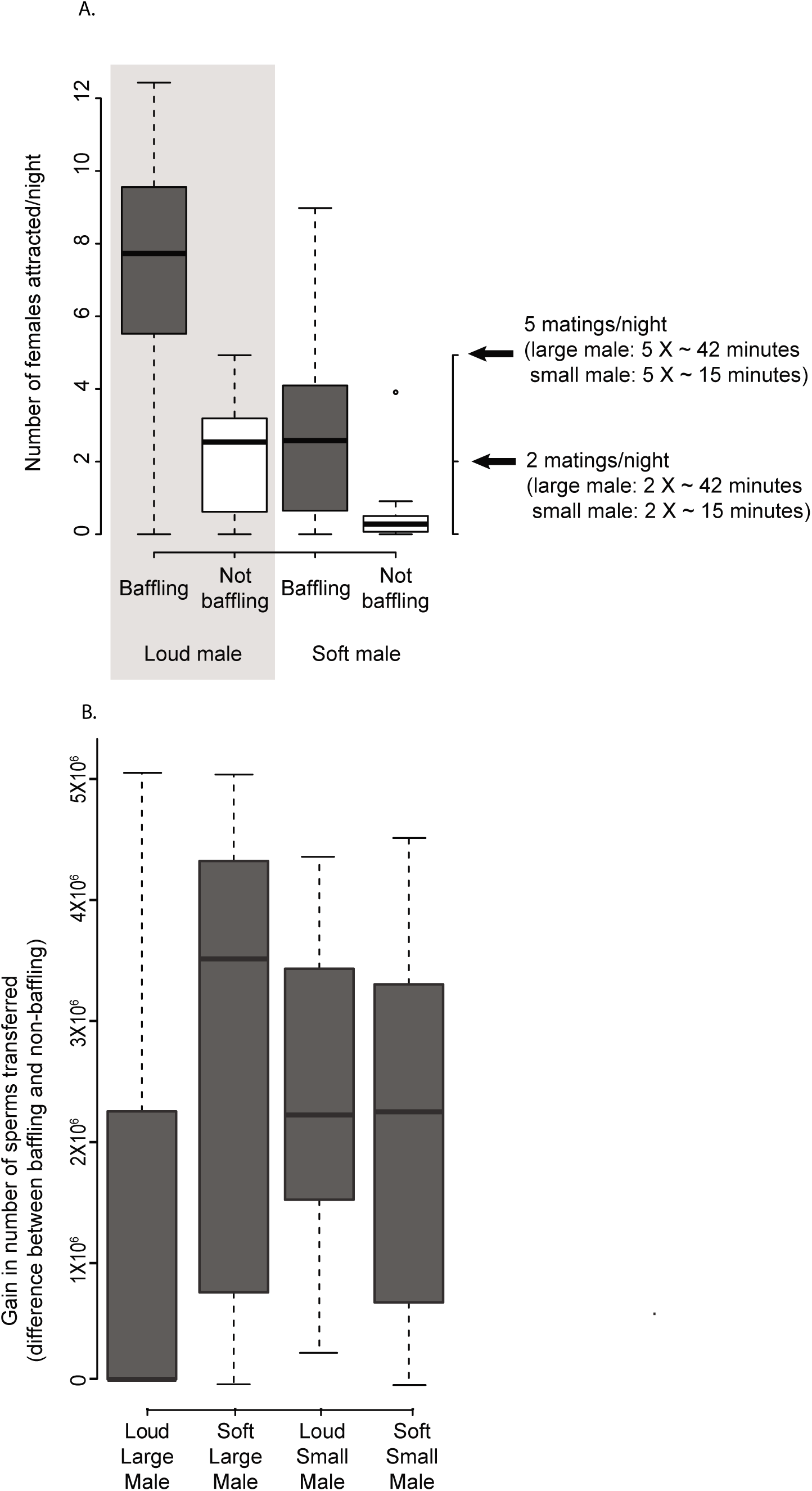
Differential benefits of baffling for males with preferred and less-preferred traits. A) Boxplots showing the number of females attracted by each type of male (loud (>66.1 dB) and soft (<61 dB)) when they were baffling (grey boxes) vs. when they were not baffling (white boxes) (each box shows the distribution across 23 choruses, simulated for 100 iterations (1 iteration = 1 calling night for an animal). Louder males attracted an optimum number of females they could mate with within a night (median of 3 females) even without baffling, whereas the softer males gained significantly when they baffled. B) Box plots showing the change in the simulated number of sperms transferred over the lifetime (LST across 100 nights) when loud and large, soft and large, loud and small, and soft and small males were baffling in comparison to when they were not-baffling. Each box shows the distribution across 22 animals across 22 choruses pooled over 100 nights in each chorus. It is evident that the less preferred males gained more by baffling. In a few choruses, the loud and large, and soft and large males did not gain by baffling – indicated by negative values in the number of sperms transferred. The large males were restricted to a maximum of 2 matings/night, whereas the small males were restricted to a maximum of 5 matings/night (see methods for more details).

We found that gain in lifetime sperm transfer (LST) when baffling was significantly different between the different groups of males (Table 1A, Fig 4B, S8). When comparing the gain in LST, we found that the loud and large males (preferred males) gained significantly less than the less preferred males (Fig 4B, Table 1A & B). The median gain in LST was highest for soft_large males (3.5×10^6^) followed by soft_small males (2.3×10^6^) and loud_small males (2.2×10^6^). The gain for the loud_large males were thee orders of magnitude less compared to other males and was negligible (3×10^3^). We found that the LST gain obtained by soft_large males were more variable (Fig 4B). These results validated our hypothesis regarding a differential gain between preferred (larger and louder) and less preferred males through baffling.

**Table 1A.**
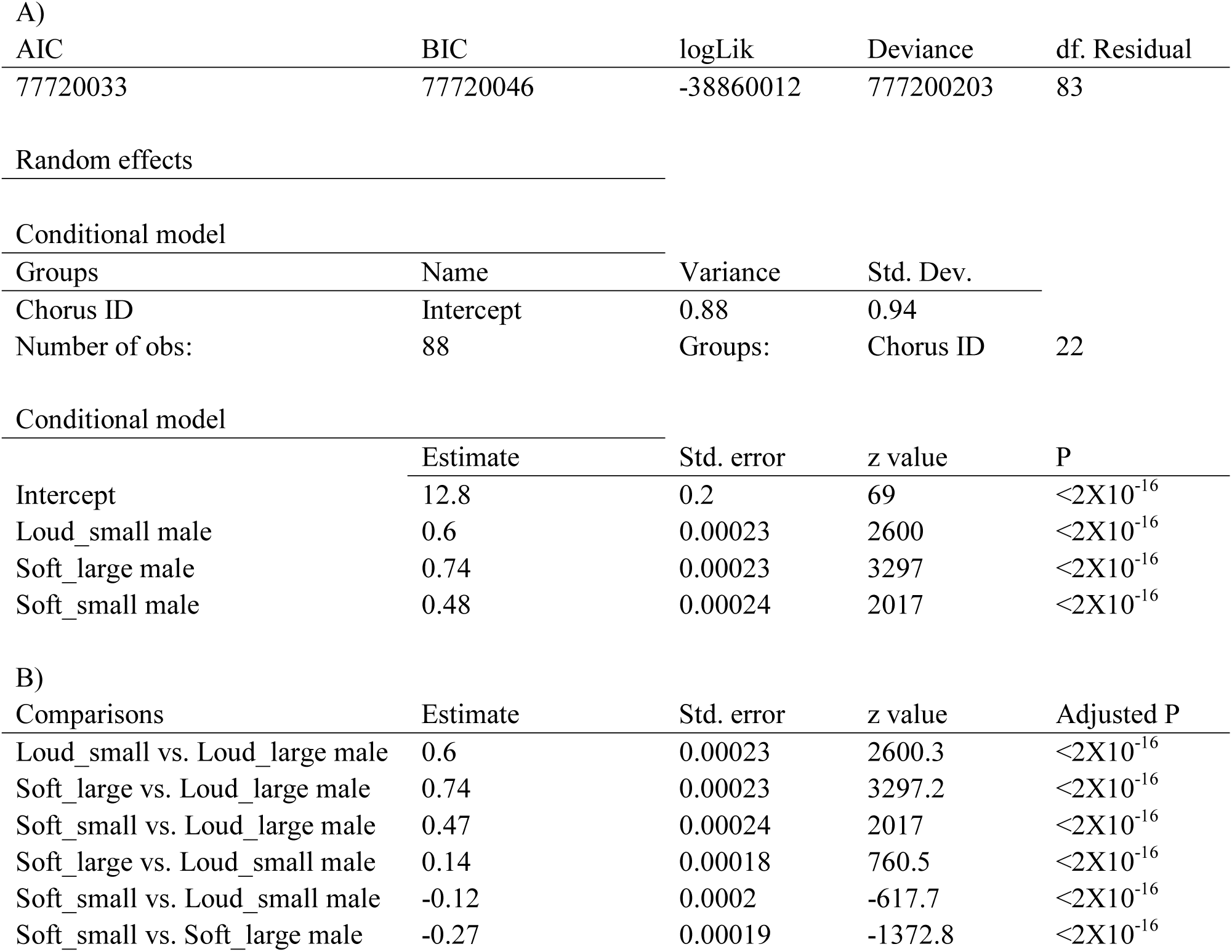
A generalized mixed model (glmm) with Poisson error showing the effect of body size and loudness on baffling gain across all pairwise combinations. B) Result of a post-hoc Tukey-test on the glmm model showing all pairwise comparisons.

## Discussion

### Baffling: a classic case of conditional alternate tactics

Earlier studies (Forrest, 1991; Mhatre, 2018; Mhatre et al., 2017; Prozesky-Schulze et al., 1975) had described baffling behaviour, optimization, and the physical principle underlying the behaviour in detail. However, none of them examined the evolutionary significance of this unique behaviour. In this study, we show that the less preferred males (smaller and with less loud calls) were more likely to use this alternative signaling tactic to gain both mating and sperm transfer benefits. Use of this alternative strategy should allow these males to reduce the inherent reproductive bias present in the species (Deb et al., 2012). Such alternative strategies based on male condition have theoretically been classified as condition-dependent strategies, where genetically monomorphic individuals choose to express a particular strategy depending on the status/state/condition of the individual (Gross, 1996; Shuster, 2010). In condition-dependent strategies, the average fitnesses of the alternative strategies are not equal, but provide higher fitness returns to the users (higher than if they were using the other dominant strategy) depending on their particular state (Badyaev, 2002; Gerhardt and Huber, 2002; Gross, 1996, 1984; Shuster, 2010; Studd and Robertson, 1985).

Gross (1996), in his seminal review of alternate strategies, laid out the critical characteristics that are essential to be examined to classify an alternative strategy as condition-dependent: a) presence of choice, b) condition/status of individuals, c) genetic monomorphism with respect to the strategy, d) fitness difference between alternatives and e) fitness advantage of the strategy. Despite decades of research, only a few studies have examined all these characteristics to classify an alternative tactic as truly condition dependent (as reviewed in Gross, 1996; Shuster, 2010; Taborsky et al., 2008). In our study, we addressed all the critical points to show that baffling is a classic case of a condition-dependent alternative strategy. A) This study in conjunction with our earlier work (Mhatre et al. (2017) established “baffling” as a facultative behaviour by showing that most males could baffle, but only a fraction of them effectively used this strategy. Our previous work (Mhatre et al., 2017) also showed that baffling propensity depended on the size of the radiator surface. These two findings, in conjunction, established that the males could choose to use this tactic. B) We show that this tactic was predominantly used by smaller males and/or those with softer calls, thus establishing that the male status/condition plays an important role in the expression of this tactic. C) Earlier work (Mhatre et al., 2017) along with our present study, also established that given an ideal surface, most males can baffle. Moreover, during our rearing of these males, we observed that newly emerged adult males kept in isolation also used this tactic (unpublished data). These indicate that baffling is unlikely to be a learned tactic, and all the males are inherently capable of expressing this behaviour. D) In our study, we provide evidence that the average fitnesses of the two alternative signaling tactics are not equal. Baffling, in general, provided higher mating and sperm transfer benefits. E) Finally, our findings indicate that the advantage gained by using this tactic was significant for the less preferred males (softer and/or smaller).

### How is the polymorphism maintained?

Maintenance of multiple signaling/mating strategies in a population has been an intriguing problem in evolutionary biology (Gadgil and Taylor, 1975; Gadgil, 1972). Alternative strategies, though they offer advantages, provide differential gains across different status/state/condition of individuals and hence might not render equal benefits to all the individuals (Gross, 1996; Shuster and Wade, 2003; Taborsky et al., 2010, 2008). Often the average fitnesses of the alternative strategies are not equal, but provide higher fitness returns to the users (higher than if they were using the other strategy) depending on their particular state (Badyaev, 2002; Bailey and Field, 2000; Reynolds and Gross, 1990; Shuster and Wade, 2003). Such differential benefits have been hypothesized to exist because mating benefits and opportunities provide diminishing returns to any dominant individual/strategy beyond a certain optimum (Real, 1980; Waltz, 1982). The main reasons behind such diminishing returns are proposed to be increasing costs to maintain dominance and lack of resource utilization (Real, 1980; Waltz, 1982). A dominant male’s reproductive success often reaches a plateau (though the number of females attracted can potentially increase linearly with male attractiveness) because it can mate only with a finite number of females within a given time (Shuster and Wade, 2003; Waltz, 1982).

Concordantly in this study, we found that though most males that baffled attracted more number of females, it was likely that the preferred males were unable to mate with all of them. The preferred males were able to attract an optimum number of females even without baffling, rendering baffling gain inconsequential. However, the benefit obtained by less preferred males from baffling was significant as they increased their mating opportunities significantly. We also hypothesized that, as preferred males were allowed to mate for extended durations (∼45 mins), their sperm transfer rate would reach a plateau (Brown, 1997; Sakaluk, 1984; Simmons et al., 2003) even without baffling. Hence, any increment in spermatophore attachment duration due to baffling will only incur minimal benefits to the preferred males. Concordant with these hypotheses we found that the reproductive gain, measured as lifetime sperm transferred (Brown, 2008, 1999, 1997; Oberhauser, 1989; Simmons, 1988), was not comparable across different males. Less preferred males gained more by baffling, whereas larger and louder (preferred) males’ gain was negligible. Moreover, we also found that the phonotactic females took significantly longer and made multiple errors while localizing the inherently loud males who were transformed into bafflers (LEL), even in our simple experimental set-up. We speculate that in field conditions, such errors will amplify causing a reduced/delayed female visitation to these males. This study thus suggests that baffling behaviour is maintained in the population along with other calling strategies because its benefits are coupled with the condition of the male (condition dependent strategy).

However, we also speculate that apart from these condition-dependent differential costs and benefits, bafflers might also face generic costs. These costs can vary from searching for ideal leaves, manufacturing cost, predation due to higher calling SPL and lesser vigilance capability (due to its typical positioning by sticking its head on the other side of the leaf). Additionally, it is known that the males of *O. henryi* provide mating incentives during copulation in the form of glandular feeding (Deb et al., 2012). Hence, we speculate that longer mating duration will increase the cost of mating per night. These generic costs should decrease the advantage of baffling and can potentially inhibit a male in favourable reproductive condition from using this strategy. It will be interesting to examine these potential costs of baffling in greater detail.

### Can baffling be termed as a cheater strategy?

Since Darwin’s proposal of the conceptual framework (Darwin, 1871) it has been widely accepted and shown that secondary sexual traits that increase individual fitness by sexual selection are inherently costly (Andersson and Iwasa, 1996; Bateson, 1983). An essential assumption of sexual selection theory has been that the marginal costs of such traits differ for different advertisers depending on their condition (Grafen, 1990; Kotiaho, 2001; Rowe and Houle, 1996). This cost essentially makes this advertising signal a reliable indicator of quality, that drives female preference. However, such differential reproductive success caused by sexual selection can lead to the evolution of “cheating” mechanisms, where individuals gain fitness benefits without paying the associated cost. There has been a long-standing debate regarding the concept and use of the word “cheat” from the evolutionary perspective. Ghoul et al. (Ghoul et al., 2014) in a recent comprehensive study suggested that in scenarios of prospective cheater behaviour it is crucial to understand the evolution of this behaviour through examining the selection acting on these cheats and the advantage the behaviour renders. In this study we found that baffling provided a two-fold reproductive gain to the males who opted for this strategy, a) increased the number of females attracted towards them and b) increased the mating duration thus allowing them to transfer more number of sperms. Females were unable to differentiate between a baffler and true loud caller (in each male size class) - during both phonotaxis and mating - and were successfully deceived. Users of this strategy, who were otherwise of a poorer condition and hence not preferred, obtained fitness benefits by bypassing/cheating the female preference for honest and reliable indicators (male size and call intensity) of male vigour. Baffling may therefore be thought of as a cheater strategy.

## Conclusion

Our previous study (Mhatre et al., 2017) had hypothesized that baffling behaviour could be a) an honest strategy exaggerating the signaling capability of the preferred males, or b) an alternative strategy allowing males in poorer condition to sound more attractive to females. In this study, we demonstrate that the less attractive smaller tree cricket males use these self-made tools (Mhatre, 2018; Mhatre et al., 2017) to deceive females into garnering more mates and longer mating duration. This study is to our knowledge the first to examine the use of self-made tools (baffles) as a condition-dependent alternative strategy in insects. Interestingly, though we understand the potential advantages of this behaviour, it remains unclear what the costs associated with it are. Finally, from a mechanistic perspective, it will be critical to understand the neuroethology of baffling that allows such tiny insects to perform such complex behaviours.

## Contributor roles

**RD:** Conceptualization, Methodology, Software, Validation, Formal analysis, Investigation, Resources, Data curation, Writing - original and final draft, review & editing, Visualization, Supervision, Project administration, Funding acquisition. **SM**: Formal analysis, Investigation, Resources, Data curation, critical manuscript review, **RB:** Conceptualization, Resources, Writing – manuscript editing and critical review, Supervision, Project administration, Funding acquisition.

## Acknowledgements

We thank R. Manjunatha, N. Ashoka, U. N. Narsimhamurthy, C. N. Chandra, and K. V. Rao for help with the collection of animals and call recordings in the field. We also thank D. Nandi, M. Bhattacharya, S. Pulla, for helpful discussions regarding the analysis and coding.

## Dataset and codes

Datasets and codes are provided as private links for the reviewing purpose. They will be made public once the manuscript is accepted.

## Materials and Methods

### The probability of finding a baffling male in the field

We localized 463 calling males across three field sites F1, F2, & F3 (see Fig. S1) over three sampling seasons. We documented the calling status, baffling or calling from other natural surfaces, for each calling male. We used these data to calculate the probability of finding a baffler in the field. During our field sampling, we had marked each male using unique colour combinations (see Deb and Balakrishnan, 2014; Deb et al., 2012); hence we knew the identity of each male. Marking allowed us to avoid pseudoreplication while sampling and recording the males.

### Who are the bafflers: preferred or non-preferred males?

#### Examining body size and calling SPL of baffling males in the field

##### Measuring body size

We localized and collected 38 non-bafflers (by random sampling) and 25 bafflers (all bafflers) from field sites F1 and F2 in separate plastic boxes (5 cm in diameter) and brought them back to the laboratory. The animals were given cold shock for 2 minutes at -4°C and then photographed along with a reference scale under a stereo zoom microscope (Leica DFC 290, Leica Microsystems GmbH, Wetzlar, Germany) with a mounted digital camera. We measured the body lengths (tip of the mandible to the anal region on the ventral side) of each animal from the images using the software ImageJ version 1.43 (National Institutes of Health, U.S.A.). The body lengths of bafflers were compared with the non-bafflers using Welch’s t-test.

##### Measuring calling SPL (loudness)

We measured the sound pressure level (SPL) (re 2 X 10^−5^ N/m^2^) (a measure of loudness) of 17 baffling and 30 non-baffling males at a distance of 20 cm (from the front, perpendicular to the wing) using a Bruel and Kjaer Sound Level Meter – Type 2231 (Bruel & Kjaer, Naerum, Denmark) with a ½□ microphone -Type 4155 (20 Hz-20 kHz) set at fast root mean square (RMS) with a flat response setting. For each SPL measurement data point, we took three SPL meter readings and calculated the average. We have reported this average value as the SPL measurement. All calling SPL measurements in the following tests were also performed using this protocol. After measuring the SPL, the baffling animal was disturbed such that it moved to a new leaf. We waited until the animal settled on the new leaf and started calling from the leaf edge. After the animal consistently called for 10 min from its new position, we measured its calling SPL. The calling SPL of bafflers when calling without a baffle was compared with that of the non-bafflers using a Welch’s t-test. Our earlier study had shown that within-night call SPL has high repeatability (Deb et al., 2012).

#### Effect of male size and call SPL on baffling probability across different leaf size classes

We conducted this experiment in the laboratory on 51 males in complete darkness during the peak calling period of *Oecanthus henryi* males (7:00 PM and 9:30 PM). The animals were chosen from three body size classes (small (<10.9 mm), medium (10.91-12.1mm), and large (>12.21 mm)), each class equally represented (17 males from each size class) (body size classes were defined based on population body size distribution (Deb et al., 2012)). As described earlier, we measured the male body sizes after cold anaesthesia one day before the experiment. Each male was kept in the experimental room 30 minutes before the experiment to acclimatize it with the room conditions. The room temperature was monitored with a Testo 110 Precision Thermometer (Testo Ltd., Hampshire, UK) and varied between 23°C-25°C.

The set-up consisted of a small *Hyptis suaveolens* (host plant) twig (15 cm in length) embedded in a thermocol piece and covered by a plastic jar. Ten such setups were prepared, and one animal was released in each and continuously observed during the entire calling period. The animals were kept in acoustic isolation. Each animal was released on four different leaf sizes over four consecutive nights using identical setups. The leaf sizes were small [3.5 (±0.2) X 2.5 (±0.11) cm], medium [4.5 (±0.2) X 3.5 (±0.1) cm], large [6.5 (±0.2) X 5 (±0.1) cm] and extra-large [11 (±0.4) X 9 (±0.2) cm] ((leaf size classes were defined based on leaf size distribution (Mhatre et al., 2017). The order of presentation of leaves was randomized (using a random number generator in R) for each animal. The animals typically started calling from the leaf edge at the beginning of the night. When an animal started calling, the plastic jar was carefully removed without interrupting the animal. As soon as the animal resumed uninterrupted calling, for at least 10 minutes, the SPL of the animal was measured at 20 cm from the front of the animal using a Bruel and Kjaer Sound Level Meter. If the animal made a baffle, the calling SPL was measured again at 20 cm from the animal. At the end of the experiment (post 9:30 PM), the animal was removed from the setup, and the baffled leaf was scanned on a flatbed scanner (HP LaserJet M10005MFP, Idaho, USA) and measured using the software ImageJ (version 1.43).

We used a generalized linear mixed model (GLMM) with binomial error structure using the R-package lme4 (Bates et al., 2013) to examine if baffling propensity was influenced by body size, without baffle calling SPL, and leaf size, with the animal ID as a random effect. As calling SPL without baffle was measured once per animal and this was a repeated measures design, we converted the SPL into a categorical variable using the calling intensity distribution of the *O. henryi* population (mean 63.8±2.55) (Deb et al., 2012). Callers whose SPL was below 63.8 were classified as soft, whereas callers whose SPL was above it were classified as loud. The basic model had all the main effects and interaction terms. We calculated analysis of deviance (Crawley, 2012; McCullagh and Nelder, 1989) for each model and examined the significance of each explanatory term. Each model was simplified stepwise by removing the non-significant terms until a saturated model was obtained. Models with acceptable fits were used for interpreting the results (model simplification). We divided the animals into size classes and examined their baffling probability based on their size, using proportion tests. Distributions of calling SPL without baffling were compared between bafflers and non-bafflers using a Welch’s t-test.

### Advantages of baffling

#### Increase in calling SPL (loudness)

We measured the call SPL of 17 baffling males in the field. Next, the baffling animal was disturbed such that it moved to a new leaf. We waited until the animal settled into the new leaf and started calling from the leaf edge. After the animal consistently called for 10 min from its new position, we measured its call SPL. The call SPLs of bafflers when calling with and without a baffle were compared using a paired t-test.

#### Experiment: Effect of male body size and leaf size on the baffling gain

In this experiment, we examined whether the gain in SPL by baffling was dependent on a) the leaf size on which the baffle was made, and b) the body size of the baffler. We expected that baffling gain should increase with an increase in the size of the radiator and a decrease in the size of the source (animal body size). We experimented on 30 males (10 large, 10 medium and 10 small males) in an anechoic chamber in complete darkness between 7:00 PM and 9:30 PM. The males were cold anaesthetized, and their body size and wing size were measured one day before the experiment. We placed each male inside the experimental room 30 minutes before the experiment to acclimatize it to the room temperature. During the experiment, the temperature of the room was monitored with a Testo 110 Precision Thermometer (Testo Ltd., Hampshire, UK). The temperature during the experiment was in the range of 23°C-25°C.

Our experimental setup consisted of a small stripped down *Hyptis suaveolens* twig, 15 cm in length, with one end embedded inside a small thermocol piece 10 cm in diameter. We attached a *Hyptis* leaf with petiole on the top of the twig and covered the junction using a wet wad of cotton to prevent dehydration. We released each animal on the leaf and covered the entire setup using a big transparent plastic jar (15 cm in diameter and 25 cm high) to prevent the animal from escaping. We prepared 4 such experimental setups. First, we released the animal on a setup consisting of a medium leaf (L X B; ∼4.5 cm X 3.5 cm) and allowed it to settle and call. Once the animal got acclimatized and started to call from the edge of the leaf, we removed the plastic jar and measured the calling SPL from the front of the animal. Next, on the same night, we moved the animal to another setup, which consisted of a small leaf (∼3.5 cm X 2.5 cm) with an artificially made baffle hole (beside the central rib) matching the size of the respective animal’s wing. Once the animal got acclimatized with the new setup and started calling from the baffle, we measured its calling SPL. Similarly, we measured the baffling SPL of the same animal from a medium (∼4.5 cm X 3.5 cm) and a large (∼6.5 cm X 5 cm) leaf set up on the same night. We randomized (using a random number generator in R) the order of presentation of the three leaves for each animal. We made the baffling holes ourselves, rather than allowing the animal to make them, as it was unlikely that the animal would make three baffle holes within a night. The experiment with an animal had to be finished within a night as calling SPL has high within night repeatability but low across-night repeatability (Deb et al., 2012). If an animal was not motivated and did not call, we observed it for 30 minutes and then replaced it with a new animal.

We calculated the gain in SPL by baffling by deducting the SPL of the animal when calling without baffle from the SPL measured while the animal was calling from baffles in small, medium and large leaves. We examined the effect of leaf size and male size on the gain in SPL via baffling using a generalized linear mixed model (GLMM) with animal identification number as the random effect. We simplified the model used the final model to interpret the results.

#### *Increase in active acoustic volume: the shape of the* active acoustic volume *of bafflers*

The active acoustic volume of a calling male is defined as the volume within which the calling SPL is greater or equal to the behaviorual hearing threshold of the female. Earlier work (Deb and Balakrishnan, 2014) showed that the behavioural hearing threshold of an *O. henryi* female is 45 dB SPL (re 2 x 10^−5^ Nm^- 2^). As the *O. henryi* males and females reside on shrubs, it is expected that the active acoustic volume of the males will be a 3-dimensional volume. To estimate the gain in the active acoustic volume through baffling, it was essential to understand the shape and volume of a caller’s active acoustic volume both when it is calling with and without a baffle. Generally, the SPL of a caller is highest at the front and back of the animal, and it attenuates towards the sides (Forrest, 1991, 1982; Mhatre et al., 2017). We designed an experiment to estimate the structure of the active acoustic volume of animals calling freely with and without baffle.

We conducted the experiment indoors in an anechoic chamber on 20 males. Our setup (Fig S3A, inspired by the set up in Forrest (1991)) consisted of a hollow aluminium tube (3 cm in diameter) of 100 cm length bent into a D-shaped half-circle of radius 25 cm with two extended arms. We covered the pipe with acoustic foam (Monarch foam India Ltd.) to reduce reverberations. We made holes of 1 cm diameter in the pipe to insert the Bruel & Kjaer microphone (½□ microphone - Type 4155, frequency response 20 Hz-20 kHz). The holes were spaced every 30° (±2°) and had screw fittings to fit the Bruel and Kjaer microphone at an angle of incidence 0° (±2°) with respect to the center. We fitted the microphone to each hole position such that the tip was placed at 20 cm from the center of the half-circle made out of the hollow aluminium tube (Fig S3A). For the experiment, we used a single Bruel & Kjaer microphone fitted to the sound level meter through a wire. The base of the aluminum semicircle (Fig S3A) was attached to a knob standing on a metal stand of height 60 cm standing on a metal plate. We fitted the knob such that the semi-circle tubing could be rotated around 360°. We also fitted the bottom plate with two protractors attached face to face, forming a circle with markings of 1° resolution (Fig S3A). We attached a laser pointer and calibrated at 0° at the bottom arm of the D-shaped tube such that the laser would point the angle of rotation on the protractor fitted at the bottom (Fig S3A).

We also prepared another stand of 90 cm length with two holding clamps of 10 cm in length. One holding clamp was at the height of 75 cm holding a thin metal plate; whereas the other clamp was above this, holding a fresh leaf with the petiole. For all trials, we fitted the leaves perpendicular to the ground such that the angle of incidence with respect to the ground was 0° (±2°). The petioles of the leaves were covered by wet cotton plugs to avoid wilting. Each night we released a single animal on a leaf attached to the setup and covered the leaf along with the holding clamp using a plastic jar (15 cm diameter and 25 cm high). The jar was slit on one end to allow the passage of the leaf holding rod, whereas the other clamp holding the plate acted as a base for the setup, preventing the animal from escaping. Each night once the animal started calling, we carefully removed the plastic jar, the plate, and the bottom holding clamp, thus leaving the animal calling from the leaf without any interference (Fig S3A). All the rods and stands were covered by acoustic foam to reduce reverberations.

We tested each animal for two consecutive nights. On one night the animal was given a small leaf (smaller than 3.5 cm X 2.5 cm) and on the other night a large leaf (larger than 6.5 cm X 5 cm). The order of presentation for each animal was randomized (using a random number generator in R). Once the animal started calling, we adjusted the height and position of the aluminium set up such that the calling animal was precisely at the center of the circle. We inserted the microphone of the sound level meter at each hole position to record the calling SPL at specific elevation points. We took 5 SPL measurements at each point and averaged the values to calculate the final SPL. Once the SPL measurements were made from all the positions of the semicircle, we rotated the setup by 30° along the azimuth, and all the measurements were retaken. Thus 84 average SPL measurements were taken around the animal forming a sphere. The measurements at 0° elevation and 180° elevation were actually at the same point for all the azimuthal angles. We used these multiple measurements from the same positions to examine if the calling SPL changed during the experiment. The entire experiment was performed at room temperature varying from 23°C-25°C.

We acclimatized each animal for 30 minutes before the start of the experiment. The experiments were conducted between 7 PM and 9:30 PM. If any animal did not call for 30 minutes, we replaced it with a new animal. We scanned the leaves used for the trials using a flatbed scanner for further analysis (length and breadth measurement).

Once all the SPL measurements were collected, using the formula for spherical spreading (Arak and Eiriksson, 1992) we converted the SPL values surrounding the animal (i.e. at source) to distance at which it attenuates to 45 dB. Earlier studies showed that *O. henryi* calls do not suffer from any excess attenuation in the field (Deb and Balakrishnan, 2014); hence, we considered only the attenuation through spherical spreading. We plotted the shape of the isobar surface in MATLAB (version 2007a) to examine the shape of the active acoustic volumes (Fig S3B, C, and D). We wrote an algorithm, based on vector multiplication, in MATLAB (see code) to assess the volume of these complex active acoustic volumes. The length of each axis (y, x, and z) depicted the length of the active acoustic volume in the direction of the axis. For example, the length of the “y” axis depicted the length of the active acoustic volume along the y-axis (Fig S3 B, C, and D).

Next, the shape and volume of these baffling active acoustic volumes were approximated with known 3D structures (such as a sphere, oblate and prolate spheroid, scalene ellipsoid) to understand the relationship between SPL and active space volume. The active space was best approximated by two identical ellipsoids touching each other along the longest radius (b) representing the body axis (y-axis) of the animal (Fig S3B, C and D, Fig S4A). The three radii of each ellipsoid were calculated from the existing 3D-active acoustic volumes (Fig. S3B, C, and D) as; 

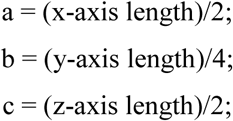

Where b = ellipsoid’s 1^st^ major axis (depicting axis y, i.e., the plane of the animal body, Fig S3B), a = ellipsoid’s 2^nd^ major axis (depicting axis x, i.e. breadth of the animal body, Fig. S3B), and c = ellipsoid’s 3^rd^ major axis (depicting axis z, i.e. height of the animal, Fig. S3B). The fit of the approximation was tested through regression analysis (Fig S4B). Post volume fitting, we calculated the relationship between the ellipsoid axis lengths (i.e., b, a, and c) and the calling SPL (ranging from 59 – 76 dB) (Fig S4C). These relationships were used for baffling active space simulations.

In order to examine the increase in active acoustic volume through baffling, we chose a range of non-baffling call SPLs, ranging from 58-68.5dB with an increment of 0.5 dB, from the known distribution of calling SPL in this species (citation?). We first mapped these SPL values onto active space volumes of non-bafflers. For non-bafflers, we considered a spherical active acoustic volume (i.e., without any attenuation on the sides). We considered spherical active space as non-baffling animals intermittently re-orient themselves across random angles within a night to advertise across 360° axes (personal observations, Rittik Deb). Following these calculations, we increased each calling SPL by 8, 10, and 12 dB SPL, respectively (typical increment in SPL when baffling). We calculated the active space volumes using the relationship between SPL and the length of the ellipsoid axis (Fig S4C). These baffling active acoustic volumes were compared with non-baffling spherical active acoustic volumes to understand gain in active acoustic volume through baffling.

#### Effect of baffling on female phonotactic preference

##### Estimating the SPL differences in the field when active spaces of multiple males overlap

Before examining female phonotactic preference based on the loudness of the call, it was essential to understand which SPL levels and SPL differences were ecologically relevant. We aimed to understand the female preference for SPL differences in scenarios where active acoustic volumes of multiple individuals overlapped. For this purpose, we used 24 natural choruses and chose those individuals that were nearest to each other (Deb and Balakrishnan, 2014), as these individuals were expected to have maximum overlap in the active acoustic volumes. We used the call SPL of each individual at the source, sound transmission pattern in the field, their location in the chorus map and nearest neighbour distances (Deb and Balakrishnan, 2014) to calculate the actual SPL levels of both the focal male and its competitor at the overlapping zone. Next, we randomly transformed the focal male into a baffler (calling SPL + 12 dB) to re-estimate the SPL at the overlapping active acoustic volumes. We plotted the SPL of the focal animal, its closest competitor, and the difference in their SPLs (Fig S5a), to understand the ecologically relevant SPLs and the difference in SPLs faced by females in natural choruses.

##### Female phonotactic preference for louder calls

###### Animals

For this experiment, we collected nymphal instars of female *O. henryi* from field site 1 (Fig S1). These female nymphs were kept individually in cylindrical plastic containers of diameter 6.5 cm and height 4.2cm with perforated lids. We provided *ad libitum* food (freshly cut apple pieces and host plant (*Hyptis suaveolens*) leaves) and water. We maintained all the animals on 12:12h light and dark cycle at room temperature, between 18°-28°C, the natural range of temperature found in the field. We used adult virgin females for the experiment within 15-21 days of their final moult.

###### Setup

We conducted all the experiments in an anechoic chamber in complete darkness. The testing arena in the chamber consisted of two loudspeakers (Creative SBS 240, Creative Technology Ltd., Singapore) standing at the height of 60 cm from the ground and separated by a distance of 120 cm. We placed stripped branches of *Hyptis suaveolens* horizontally between the two playback speakers as a bridge, supported at right angles by a similar vertical branch placed equidistantly from the speakers in the same horizontal line. This essentially formed a T-junction with the speakers at the two extremities (Fig. S5B). We carried out the trials between 7:00 to 9:30 PM, at an artificially maintained mean temperature of 25°C (24.5-25.5°C). The females were acclimatized in the test chamber at 25°C for 1.5 hours (1 hour light, 30 mins in darkness) before the experiments. We conducted two-choice playback experiments using combinations of calling SPL generated from the field data (Fig S5A) with both speakers broadcasting simultaneously. For each trial, we released an individual female at the base of the central vertical branch after starting the playback. The female had a choice to walk towards either of the speakers. We set a cut-off of 120 s for the females to show any response towards a speaker (Deb and Balakrishnan, 2014; Deb et al., 2012). We recorded the movements of the animals using an IR-sensitive video camera (Sony DCR-TRV 17E, Sony Corporation, Tokyo, Japan).

###### Experimental design

In our experimental design, in each trial, both the speakers were playing back naturally recorded identical *O. henryi* male call (with temporal settings specific to 25°C (Metrani and Balakrishnan, 2005)), only varying in SPL and phase relationship. The preference of a female for a given calling SPL was determined by its phonotactic response towards the speaker. We set the absolute SPL as 50 dB (re 2 x ^10-5^ N/m^2^ based on field recordings, Fig S5A). We chose 3 combinations of SPL pairs for playback: 50dB - 53dB, 50dB - 56dB, 50dB -59dB, with a difference of 3dB, 6 dB, and 9 dB, respectively. The SPL of the acoustic stimulus from each speaker was adjusted at the T-junction before each trial using a sound level meter. We carried out two control trials for every experimental session; a) a silent trial where no acoustic stimulus was presented and b) where both the speakers were simultaneously played at the base SPL of 50 dB. The silent control determined the probability of the female to respond in the absence of any acoustic stimuli. The 50dB - 50dB (0 dB SPL difference) control checked for response biases. We randomized (using a random number generator in R) the directionality of the louder speaker for each test trial for each animal. We tested 2-3 animals per night to reach sample sizes of 16 (50-53 dB), 20 (50-56 dB) and 21 (50-59 dB) across the trials. If an animal reached a speaker broadcasting a particular call within a cut-off period of 120 s, only then we considered it as a response (hence unequal sample size across trials). If it failed to reach a speaker within the cut-off period, we considered it as a “no response.” The cut-off was established based on previously published data (Deb et al., 2012; Mhatre et al., 2011). We carried out 5 trials/night, 3 test + 2 control, for each female. We maintained a gap of 30 minutes between successive trials for an individual animal to eliminate any effect of the previous trial on the response. We randomized (using a random number generator in R) the order of all the test and control trials for each female. We performed a Chi-square test to examine if the females preferred the louder calls in each trial. We also adjusted our p-values for multiple comparisons using false discovery rate correction.

#### Effect of baffling on male mating opportunities

We tested the increase in mating opportunities (proportion of females attracted) obtained by baffling males using a simulation. For the simulation, 24 natural chorus spatial maps along with measured male calling SPLs were used (Deb and Balakrishnan, 2014). To measure the gain in mating opportunities via baffling, we simulated each chorus twice, **Scenario 1:** without any baffler in the chorus, and **Scenario 2:** with a randomly chosen male transformed into a baffler by increasing its calling SPL (between 8-12 dB) and typical baffling active space (two connected ellipsoids fixed at a fixed angle). We kept the angle fixed as a baffler’s active acoustic shape is governed by the angle of the leaf from which it is calling, which remains constant within a night. The angle of the leaf was randomly changed every night (considering a new baffle). For all the non-bafflers, we simulated freely rotatable (360°) active acoustic volume around the center of the axis (the animal position) by rotating their calling angle after a random interval ranging between 15-40 minutes (range obtained from personal observations made by Rittik Deb). The radius of the active acoustic volume was calculated using the relationship between the calling SPL and ellipsoid radius (Fig S4C).

For the simulation, we considered only the females that occurred inside the active acoustic volume of a calling male. We assumed that females outside any male’s active acoustic volume would show a random walk (Gerhardt and Huber, 2002) as it has no directional cue and would only be relevant once it enters the active acoustic volume of a male. This also meant that the larger the active acoustic volume, and more omnidirectional it is, higher will be the probability of a female landing inside it.

For the simulation, across both scenarios (1 and 2), we kept the calling male to female sex ratio for each chorus equal (1:1), i.e., if a chorus had 10 males calling in it, 10 females were assigned to random positions (1cm pixel size) in that chorus for that night (i.e., that simulation run). If a female was positioned inside the active acoustic volume of multiple calling males, then we calculated the SPL (dB) of all the males at that point. The decision to choose a male was decided using the a) the SPL difference between the multiple males at that point, and b) outcome of female preference experiment for louder calls. If the males matched in their calling SPL at the female position, then we randomly assigned the female to one of the males. If a female’s position was in the active acoustic volume of one male, we assigned the female to that male. This simulation was run until the number of females assigned was equal to the number of males present in the chorus. At the end of the simulation run, we recorded the number of females obtained by each male. This process was iterated for 100 times for each chorus, representing 100 nights for each male. We ran it for 100 nights to calculate an estimate of the lifetime mating success for each male (mean male lifespan 100 days in the laboratory) in each chorus (considering the rarity of mate rejection in *O. henryi* post localization, Rittik Deb personal observation). We transformed the number of females obtained by each male into the proportion of females obtained, by dividing the number of females obtained by each male in a chorus by the total number of females assigned to that chorus.

At the end of 100 iterations, the proportion of females obtained by a baffler male while baffling and while not baffling was compared across all the 24 choruses using the pairwise-randomization test. The proportion of females obtained by each male was plotted against their call SPLs for both the chorus scenarios (scenario 1 (all non-baffler) and 2 (only 1 baffler, rest non-baffler)). We performed a generalized linear model with a binomial error, where the proportion of females obtained was the response variable (taken as failures and successes in obtaining a female), and the calling SPL without baffle was the predictor (scenario 1). This was performed to examine if the number of females obtained without baffling increases with calling SPL (scenario 1). This same model was rerun for scenario 2 – i.e. when one male in the chorus was baffling. We also calculated Z-scores for all the males based on the proportion of females attracted and plotted them against the calling SPLs.

#### Effect of baffling on female phonotaxis and mating behaviour

##### Animals

For the experiment, we used adult virgin females (within 15-21 days of their final moult). These females were collected as nymphs from the field site 1 (Fig S1) and maintained in the laboratory until adult eclosion (as stated earlier). We used freshly caught males from the nearby field site (field site 2: 100m away from the field station) for the experiment. Each night we located calling males and recorded their calling SPL in the field. Following this, we captured the males, along with the leaf from which they were calling, inside cylindrical plastic containers (6.5 cm diameter X 4.2 cm height) and brought them back to the field station immediately. Each animal was acclimatized to the room conditions in the field station under complete darkness for 30 minutes before the start of the experiment.

##### Setup

We conducted the experiment in the field station (near sampling site F2, and F3) in a dark anechoic chamber (10 X 10 ft). We made the room anechoic by covering the walls and grounds of the entire room using 150 mm acoustic foam sheets (Monarch Foams India Ltd., Bangalore). The test setup was made of *Hyptis suaveolens* branches and was almost identical to our earlier phonotaxis experimental setup (see section: Female phonotactic preference for louder calls; Fig S5C). We modified our earlier set up such that it allowed a phonotactic female to mate with a male at the end of her phonotaxis. For this purpose, we prepared a Petri-dish (10 cm diameter) as a mating chamber (as used in Deb et al. (2012)) and kept it at the very end of the arm leading to the playback speaker. We covered the base of the Petri-dish with fresh *Hyptis suaveolens* leaves to make the setup more natural. For all the trials, we released the females at the base of the central stick and recorded the animal movement using an IR sensitive camera. The setup consisted of two speakers (X-mini speakers, Xmini, Japan) connected to a laptop (Vostro 1400, Dell, Texas, USA) for playback. We performed a no-choice experiment where only one of the two speakers was playing. The animal had the choice of approaching the playback speaker, remain at the decision point or move away from the playback speaker towards the silent speaker. Once the female had reached the Petri dish, we used another Petri-dish (of exact similar dimension) as a cover to prevent the female from escaping. We released the male whose call SPL was being played into the Petri-dish. We stopped the playback as soon as the male and the female antennated, simulating a natural phonotaxis and mating scenario. We conducted the experiments at room temperature varying between 26.5°C-27.6°C. The experiments were conducted between 7:30 PM to 9:30 PM.

##### Experimental design

We aimed to examine a) if the females preferred louder males both during phonotaxis (i.e., faster response) and mating (i.e., longer spermatophore attachment duration), and b) if the females could differentiate between a naturally loud male and an artificially loud (baffling) male. The stimuli played back consisted of *O. henryi* calling song with the temporal and spectral features appropriate for the mean temperature of the room (27°C) (Deb et al., 2012; Metrani and Balakrishnan, 2005) but varied in calling SPL. The SPL of the playback depended on a) the focal male’s calling SPL in the field on that night, and b) whether it was a baffling or a non-baffling treatment. We adjusted the SPL of the playback by measuring the playback SPL at the decision junction (middle of the setup) using a sound level meter.

Each night during male collection, we visited the field site 1 and localized calling males and measured their calling SPL. We caught males who fell in the category of soft (< Mean [63.8dB] -1SD [2.5] = < 61 dB) or loud (Mean+1SD) (>66 dB) and brought them back to the field station for the experiment (SPL distribution 63.8±2.5 dB, (Deb and Balakrishnan, 2014). We caught a total of 60 such males for the experiment out of which 40 were soft callers, and 20 were loud callers. Out of these 40 soft callers, we selected 20 random males (randomized using a random number generator in R) and artificially ‘transformed’ them into ‘bafflers’ from the female perspective. This was achieved by increasing their call SPL during playback (SL: soft transformed to loud). The increment in SPL was picked between 8-12 dB (range of gain in SPL through baffling) using a random number generator. For the other 20 soft males, the playback was performed at their original SPL (SS: soft remained soft). Similarly, for the loud callers, we played back the call of 10 males at their original field recorded SPL (LL: loud remained loud), and we ‘transformed’ 10 males into ‘bafflers’ (LEL: loud transformed to extra loud). For each male, a single female was tested for phonotaxis and mating. For each male and female pair, we tested the female for two phonotactic treatments, 1) a silent control: where both the speakers were silent, 2) playback treatment: where only a single speaker played back. We randomized (using a random number generator in R) the direction of the playback speaker across the males to avoid any directional biases in the female response. Once the female reached the Petri-dish, the respective male, whose calling SPL was played back, was released. Once the individuals antennated with each other, we stopped the playback and recorded the entire mating session. If a female did not show phonotaxis during the playback treatment, we replaced the female with a new animal from our stock. After the experiment, we measured the body lengths of the males under a microscope. We sorted the males to pre-defined body size classes (S-small, M-medium, L-large) (Deb et al., 2012) for further analysis. In our experiment, we had two treatment factors a) playback with 4 levels and b) body size with 3 levels. We divided the whole sample subset into 12 combinations, such as S_SS (Small and soft male remained soft), S_SL (Small and soft male transformed to a baffler), S_LL (Small and loud male remained loud) and S_LEL (Small and loud male transformed to a baffler), M_SS, M_SL, M_LL, M_LEL (for medium-sized males), L_SS, L_SL, L_LL, L_LEL (for large-sized males).

##### Analysis

We calculated the latency of response by measuring the time taken by each female to move from the release point to the Petri-dish in front of the playback speaker. We compared the latencies across the four playback treatments (SS, SL, LL, LEL) using pairwise ANOVA and post hoc Tukey tests. We also measured the duration of spermatophore attached to the female body (Spermatophore attachment duration; SPAD) as a proxy for female mating preference for a male (Deb et al., 2012). We performed an ANOVA to examine the effect of body size and treatment on the SPAD. We hypothesized that if the females could identify the genuine loud caller either through call or during mating, they will discriminate between LL (true loud) and SL (baffling loud) males. On the contrary, if baffling influenced female preference, the SL (baffling loud) males will have higher preferences than SS (genuine soft) males.

#### Why don’t larger and louder males baffle as much as smaller and softer males?

##### Is the gain in SPAD due to baffling comparable between small_soft and large_soft males?

For this analysis, we first calculated the gain in SPAD for small_soft males when they were transformed into bafflers (i.e., S_SL – S_SS) using the result of the previous experiment. Similarly, we calculated the gain in SPAD for large_soft males when they were transformed into bafflers (i.e., L_SL - L_SS). The purpose of this analysis was to understand whether less preferred small and soft males gained more by baffling. As our earlier experiment was performed on independent individual animals, we calculated all possible differences in SPAD between each S_SL and each S_SS data points. From this, we generated the distribution of the differences for small males. Similarly, we also prepared the distribution of the difference between L_SL and L_SS treatment. These two distributions were compared using the Mann-Whitney-U test to examine whether smaller males gained more in SPAD by transforming themselves into bafflers.

##### Is the gain of baffling (number of females attracted and amount of sperm transferred) comparable between preferred (large and loud) and less preferred (small and soft) males?

We compared the gain of baffling (both number of females attracted and the amount of sperm transferred) between preferred and less preferred males using a simulation framework. For this purpose, we simulated each of our 23 natural choruses 4 times. In the first simulation, we simulated to calculate the number and proportion of females (over its life-time, i.e., 100 nights) attracted by a randomly chosen loud male (SPL >66.1 dB) in each chorus. In the 2^nd^ simulation, we transformed these loud males into bafflers (without altering their position in the chorus) and re-calculated the number and proportion of females they attracted over its lifetime. The loud males were transformed into bafflers by a) increasing their SPL (using a randomly chosen value between 8-12 dB), and b) restricting their advertisement direction (explained in earlier simulation). Similarly, in the 3^rd^ round of the simulation, we chose a soft male in each chorus and examined the number and proportion of females they attracted over their lifetime. In our 4^th^ round of the simulation, we transformed these soft males into bafflers and re-examined the number and proportion of females they attracted over their lifetime (100 nights). We then divided the number of females attracted over their lifetime by 100 (i.e., 100 nights of calling) to calculate an average number of females these males attracted/night by calling with or without baffle. We plotted these values for loud_baffler, loud_non_baffler, soft_baffler, and soft_non_baffler across 23 choruses and compared the medians.

###### Sperm transfer function

We calculated the sperm transfer function, i.e. the number of sperms that get transferred as a function of time, using the data from a conspecific species *Oecanthus nigricornis* (Brown, 1997). This species of *Oecanthus* is comparable to the *O. henryi* in mating durations (Brown, 1997), hence we considered it to provide a realistic estimate of sperm transfer for our study. Using the data collected from this study (from fig 2, and main text), we calculated the total number of sperm that is retained in spermatophore (Table S3). We considered the upper standard error of the data (fig 2, Brown, 1997) as a representative sperm count for large males, whereas the lower bound was considered as the sperm count of a small male at each time point (Brown, 1997). We calculated the number of sperms that were retained in the spermatophore (and converting to the original value, i.e. without dilution) and fitted exponential models. The fitted exponential models to obtain best-fit lines for this data (0-30 minutes of spermatophore attachment duration) for each body size are, large male: y=29988*exp^(−0.0006*x)^, R^2^=0.99, medium male: y=24442*e^(−0.0007*x)^, R^2^=0.99, small male: y=19109*e^(−0.0009*x)^, R^2^=0.99, 

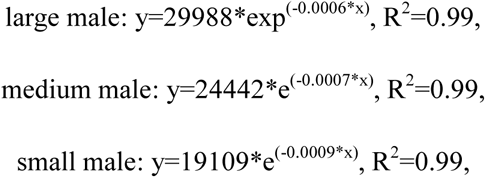

where y is the number of sperms remained in spermatophore, and x is the spermatophore attachment duration in minutes (Fig S7A). Using these fitted lines, we extrapolated the relationship between sperm remained in spermatophore for 45 min and 60 mins spermatophore attachment durations (Table S3). Next, we deducted each of these values from the total sperm remaining in the spermatophore at 0 minute, to plot sperm transfer as a function of time (Fig S7B). We fitted best-fit lines to this plot for small, medium and large males to calculate the sperm transfer function for all body sizes large male: y= -0.0023x^2^ + 15.57x, R^2^ = 0.99, medium male: y = -0.0018x^2^ + 12.468x, R^2^ = 0.99, small male: y = -0.0013x^2^ + 9.3996x, R^2^ = 0.99, 

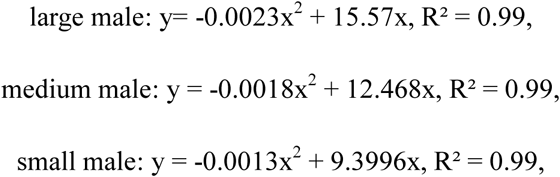

where y is the number of sperms transferred, and x is time (Fig S7B).

We multiplied these sperm transfer functions (for respective body size males), with the respective mating durations (obtained from our earlier experiment, Fig 3B) to calculate the amount of sperm that a male could transfer for a given mating depending on its calling SPL and body size. Next, we multiplied these with number of mates a male can mate within its lifetime (i.e. simulation duration of 100 nights) across each chorus. We calculated different re-mating scenarios/night, where a male could mate with 1-5 mates/night. We limited the maximum number of matings for a large male to 2 mate/night (described above) and for a small male to 5 mate/night. We deducted the lifetime sperm transfer of a male when it was not baffling from the lifetime sperm transfer when it was not baffling. This value was considered as the gain in sperm transfer by baffling for each size and loudness class of males (Fig 4B, and S8).

## Supplementary file

We have a single supplementary file with all the supplementary tables and figures with legends embedded in it. The file is named “Supplementary_eLife_Research_Advance_Deb et al.”.

